# Parent-offspring inference in inbred populations

**DOI:** 10.1101/2021.11.23.469656

**Authors:** Jan-Niklas Runge, Barbara König, Anna K. Lindholm, Andres Bendesky

## Abstract

Genealogical relationships are fundamental components of genetic studies. However, it is often challenging to infer correct and complete pedigrees even when genome-wide information is available. For example, inbreeding can obfuscate genetic differences between individuals, making it difficult to even distinguish first-degree relatives such as parent-offspring from full siblings. Similarly, genotyping errors can interfere with the detection of genetic similarity between parents and their offspring. Inbreeding is common in natural, domesticated, and experimental populations and genotyping of these populations often has more errors than in human datasets, so efficient methods for building pedigrees under these conditions are necessary. Here, we present a new method for parent-offspring inference in inbred pedigrees called *SPORE* (Specific Parent-Offspring Relationship Estimation). *SPORE* is vastly superior to existing pedigree-inference methods at detecting parent-offspring relationships, in particular when inbreeding is high or in the presence of genotyping errors, or both. *SPORE* therefore fills an important void in the arsenal of pedigree inference tools.

**Author Summary:** Knowing the genealogical relationships among individuals is critical for genetic analyses, such as for identifying the mutations that cause diseases or that contribute to valuable agricultural traits such as milk production. Although many tools exist for establishing pedigrees using genetic information, these tools fail when individuals are highly inbred, such as in domesticated animals, or in groups of people in which consanguineous matings are common. Furthermore, existing tools do not work well when genetic information has errors at levels observed in modern datasets. Here, we present a novel approach to solve these problems. Our method is significantly more accurate than existing tools and more tolerant of errors in the genetic data. We expect that our method, which is simple to use and computationally efficient, will be a useful tool in a diversity of settings, from the studies of human and natural populations, to agricultural and experimental settings.

## Introduction

Genealogical relationships among individuals are critical for many genetic analyses. For example, in trait-mapping studies based on linkage [1] and transmission disequilibrium analyses (e.g. [2]) as well as in sibling-based genome-wide association studies [3], the transmission of traits and alleles is analyzed within pedigrees to control for environmental similarities and to avoid confounders from population structure. Pedigrees can also be fit into statistical models to estimate the heritabilities of traits [4,5]. Removing close relatives from analyses can further be important to fulfill analytical assumptions in population genetics [6]. Pedigrees are also useful to study the rate of *de novo* mutations and their impact on traits [7]. Furthermore, pedigrees themselves can be of considerable interest, for example when analyzing differences in lifetime reproductive success [8] or assortative mating [9]. Thus, accurate and complete pedigree relationships are important to maximize power and accuracy of genetic analyses and for studies of population biology.

Deriving complete and correct pedigrees can be challenging. In monitored wild populations, researchers may use a combination of observed data, such as birth and death dates, coupled with sparse genotypes produced by technologies such as microsatellites, and employ software tools such as *COLONY* [6] to build pedigrees. This approach is usually accurate, but has two major drawbacks. First, documenting birth and death dates (to exclude impossible parent-offspring matches) can be difficult in some species and contexts. Second, it is impractical to genotype large numbers of microsatellites, decreasing the power to infer genealogical relationships in populations with low genetic diversity.

Current genome-wide methods for pedigree reconstruction are often developed and optimized for outbred human populations and thus compatibility with considerable levels of inbreeding is not the main concern (e.g. [10]). However, some human populations and individuals have high levels of inbreeding, with up to 20% of their genome being homozygous [11,12] (**Figure 1 A**). Considerable inbreeding is also present in populations of other species, including livestock ([13,14], and Methods), small populations of wild animals such as wolves [15] and Devils Hole pupfish [16], and even in populations with large census sizes such as house mice (**Figure 1 A**). Inbreeding violates the assumptions of many genealogical-relationship inferring software, yet the performance of these tools in the presence of inbreeding has not been thoroughly evaluated. Inbreeding homogenizes genomes within populations, obfuscating differences in the transmission and recombination of genomes between different types of relatives. For example, the fractions of their genomes two individuals share identical-by-descent (“IBD,” inherited through a recent common ancestor), in only one of the two chromosomes in a pair (“IBD1”) or in both chromosomes in a pair (“IBD2”), differ between parent-offspring and full-sibling relationships. Hence, these metrics are often used to distinguish such relationships [17–19]. However, when parents are genetically related, the fraction of the genome that is IBD2 with their offspring is higher than in outbred populations and is thus more similar to full-sibling relationships (**Figure 1 B**). Thus, inbreeding could hinder the accuracy of existing pedigree-reconstruction tools.

**Figure 1:**
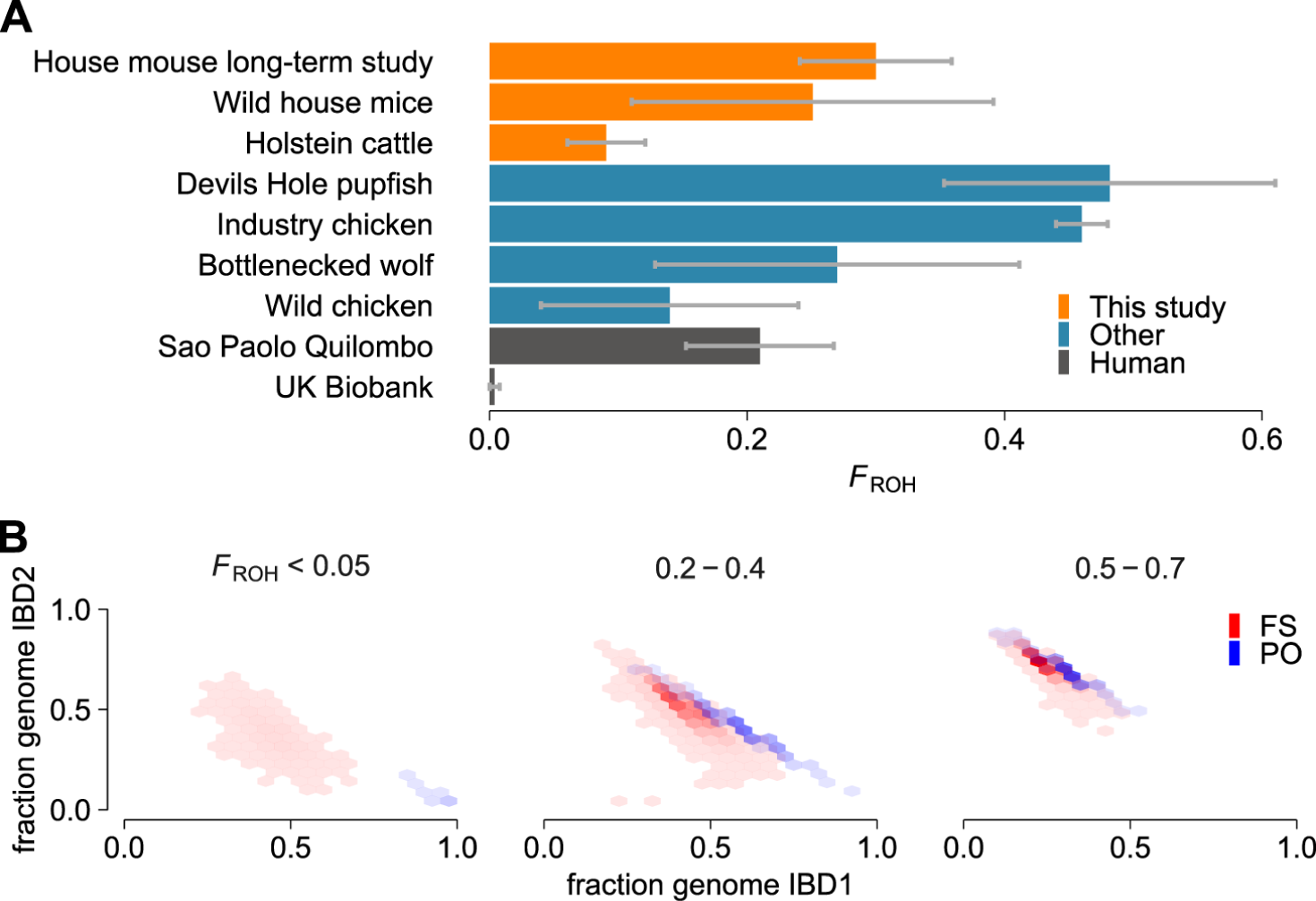
Extent of inbreeding in multiple populations and its influence on metrics used to infer relationships. **A)** Overview of levels of inbreeding, measured as *F*_ROH_ (see Methods), in the populations studied here (orange) and in other non-human (blue) and human (grey) populations [14–16,25,26]. Bars indicate average, error bars indicate the standard deviation. **B)** Joint distribution of the fractions of genomes that are IBD1 and IBD2 in parent-offspring (PO, blue) and full sibling (FS, red) relationships. Each plot corresponds to different *F*_ROH_ levels. Data is based on the simulated pedigrees with genotyping errors (see Methods).

Typical genotyping error levels in human studies can also have a large impact on pedigree inference [20,21] and these errors might be more prevalent in less well-studied species and populations [22–24]. Genotyping errors can result in parent-offspring relationships appearing as less related than expected, obfuscating the distinction between parent-offspring and other types of relationships. Furthermore, these errors can interrupt inferred IBD segments, limiting the utility of segment number and length on genealogical inference [21].

Here we present a new method, which we call *SPORE* (Specific Parent-Offspring Relationship Estimation), that is robust to high levels of inbreeding and errors in genotyping. It is designed to use genome-wide genotypes from a variety of sources, such as those derived from high-coverage sequencing, imputation, or genotyping arrays. Instead of the commonly used IBD1 and IBD2, our method relies on related allele-sharing metrics that have large expected differences between parent-offspring and other types of relationships, even in highly inbred individuals. We show that *SPORE* is more sensitive and accurate than other commonly used methods that are not designed for inbred populations and that *SPORE* is more tolerant to genotyping errors, making it a robust tool for detecting relatives and building pedigrees using genetic data.

## *SPORE* (Specific Parent-Offspring Relationship Estimation)

We developed *SPORE* as a robust method for parent-offspring inference and pedigree building in inbred populations and to be tolerant to genotyping errors. The main differences with existing methods are 1) the focus on variables that are expected to be 0 in parent-offspring relationships but higher in other relationships (with the exception of monozygotic twins) and 2) the automatic adjustment of thresholds below which a relationship is classified as parent-offspring. This contrasts with other methods’ use of fixed thresholds derived from theory, which can be violated by inbreeding and genotyping errors.

### Variables used to infer parent-offspring relationships

To infer parent-offspring relationships from genome-wide genotypes of a set of individuals, we use three variables: the proportion of the genome that is not IBD (“IBD0”), the variation (interquartile range; IQR) in that proportion among chromosomes (“IBD0 IQR”), and the proportion of loci that carry opposite homozygous genotypes (“homozygous mismatches, HM”). In outbred families, these variables have an expected value of zero in parent-offspring relationships (and in monozygotic twins), but higher than zero in full siblings and other types of relationships (**Figure 2**). IBD0 is expected to be invariably zero in all autosomes of parent-offspring pairs, yielding a total IBD0 of zero. In contrast, in full siblings on average ¼ of their autosomes are IBD2, ½ IBD1, and ¼ IBD0. However, because of recombination and the independent assortment of chromosomes, this extent of sharing varies between chromosomes, and IBD IQR will be greater than zero. Notably, interchromosomal variation in IBD0 has not previously been leveraged as a metric to infer genealogical relatives [10].

**Figure 2:**
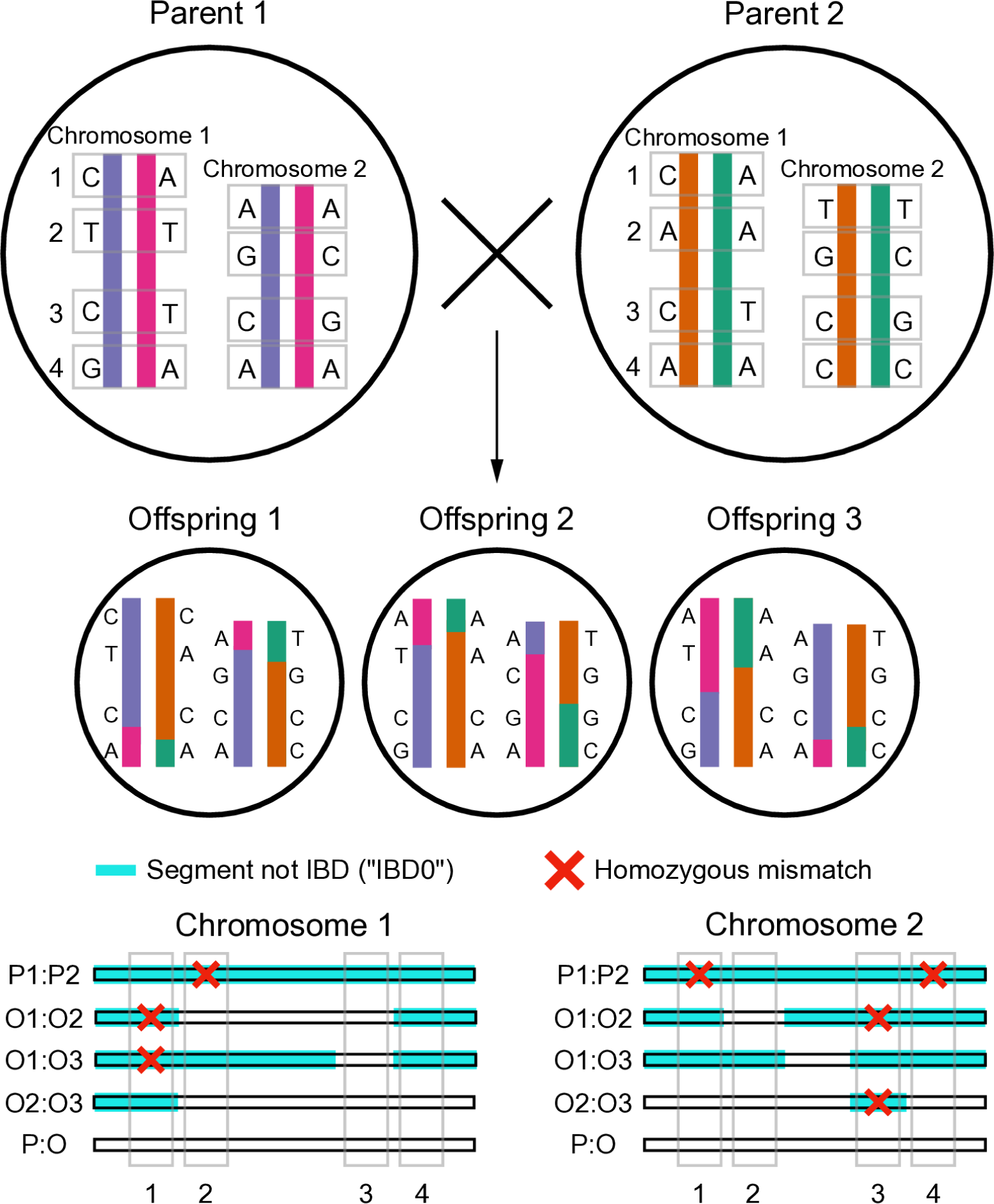
Schematic of metrics used by *SPORE* to infer parent-offspring relationships. Example of two unrelated parents who produce three offspring. Colored chromosomes denote four different haplotypes (blue, pink, orange, and green), which are recombined and transmitted to three offspring (O1 to O3). DNA bases are shown at four loci for each chromatid of two chromosome pairs. Bottom shows a comparison of the fraction of the genome that is not identical by descent (“IBD0,” turquoise), the variation in IBD0 between chromosomes, and homozygous mismatches (red crosses) across pairs of individuals. Note that full-sibling (O:O) relationships differ in all three of these values from parent-offspring relationships (P:O), which are consistently 0: 0% of the genome IBD0, 0 variation in IBD0 (“IBD0 IQR”), 0 homozygous mismatches.

Whereas IBD0, IBD0 IQR, and HM have expected values of zero in outbred parent-offspring relationships and higher than zero in other types of relationships, these metrics also move towards zero in full-siblings and other relationships under increased inbreeding (**Figure 3 A**). Furthermore, genotyping errors can result in non-zero values for these metrics in parent-offspring relationships even in outbred individuals. Hence, by contrast to other methods developed for outbred individuals (e.g. [27,28]), we do not rely on hard-set thresholds below which we consider relationships to be parent-offspring. Instead, we use the information in the dataset and a researcher-derived input variable — the assumed average parent-offspring relationships per individual (APO) — to automatically define those thresholds. Furthermore, we make use of a powerful metric, the fraction of loci that carry genotypes that could not have been transmitted from a given set of two parents to their offspring in the absence of germline or embryonic *de novo* mutations (e.g. an A/A offspring with A/G and G/G putative parents), called “Mendelian trio errors,” to determine correct father-mother-offspring trios. This metric performs well even under high inbreeding (**Figure 3 B**), but is limited to cases where full trios are present in a set of individuals genotyped densely.

**Figure 3:**
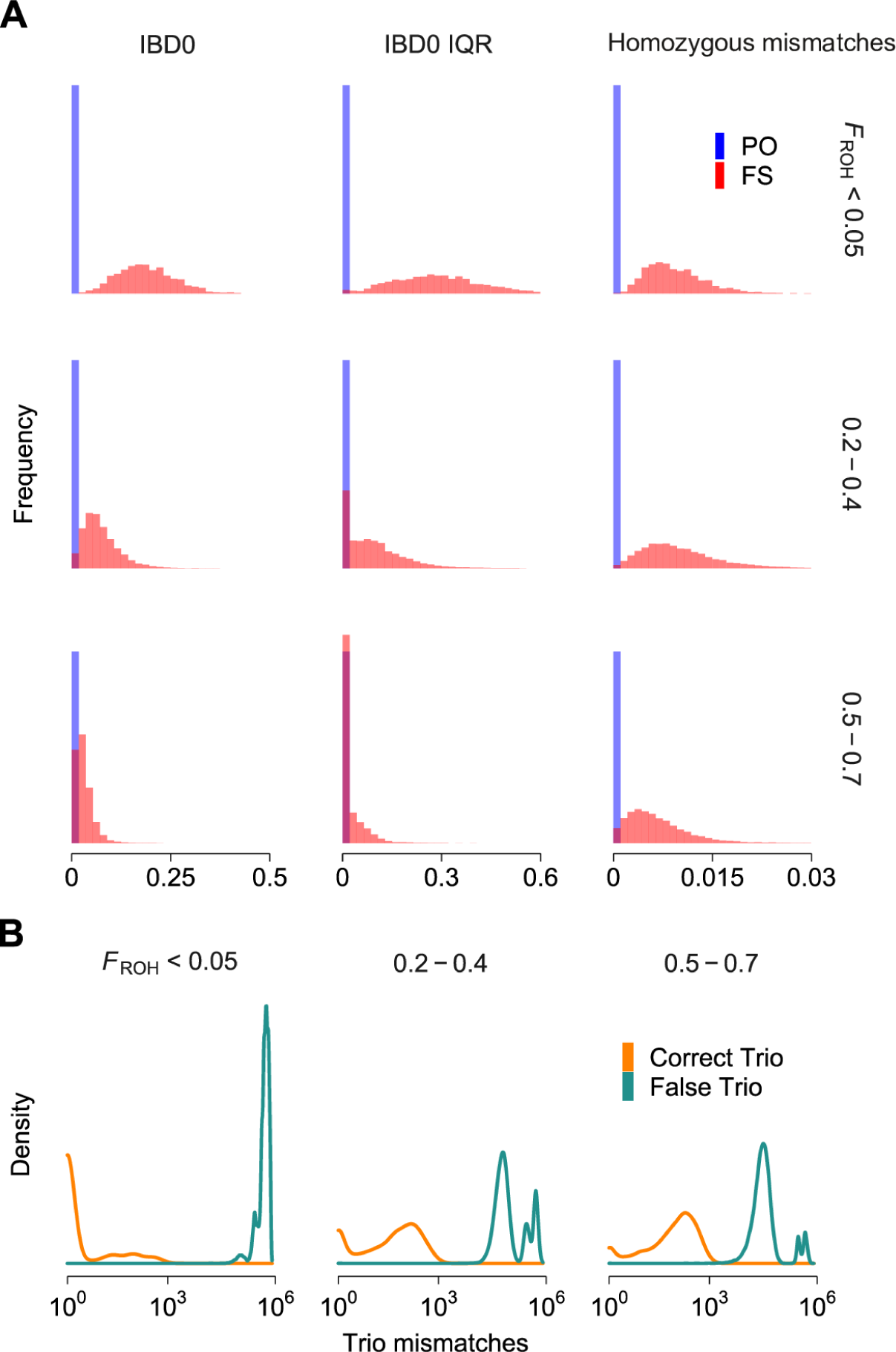
Overview of metrics used by *SPORE* to infer parent-offspring relationships. **A)** Distributions of the variables that *SPORE* uses to distinguish parent-offspring (blue) from full sibling (red) relationships, across levels of inbreeding *F*_ROH_: the fraction of the genome that is not identical by descent (“IBD0”), the inter-chromosomal variation in IBD0 (measured as interquartile range) “IBD0 IQR,” and the fraction of homozygous mismatches. **B)** Distributions of number of incompatible genotypes in true trios (orange) and in putative, but with at least one incorrect, parents (green) across levels of inbreeding *F*_ROH_. Data in **A** and **B** are based on the simulated pedigrees with genotyping errors (see Methods).

### Algorithm

*SPORE* is divided into three phases: 1) Detection of putative parent-offspring (PO) relationships (**Figure 4, 1-3**), 2) inference of father-mother-offspring trios within the putative PO relationships (**Figure 4, 4-7**), and 3) inference of PO relationships in the remaining putative PO relationships that could not be assigned to trios (**Figure 4, 8-10**).

**Figure 4:**
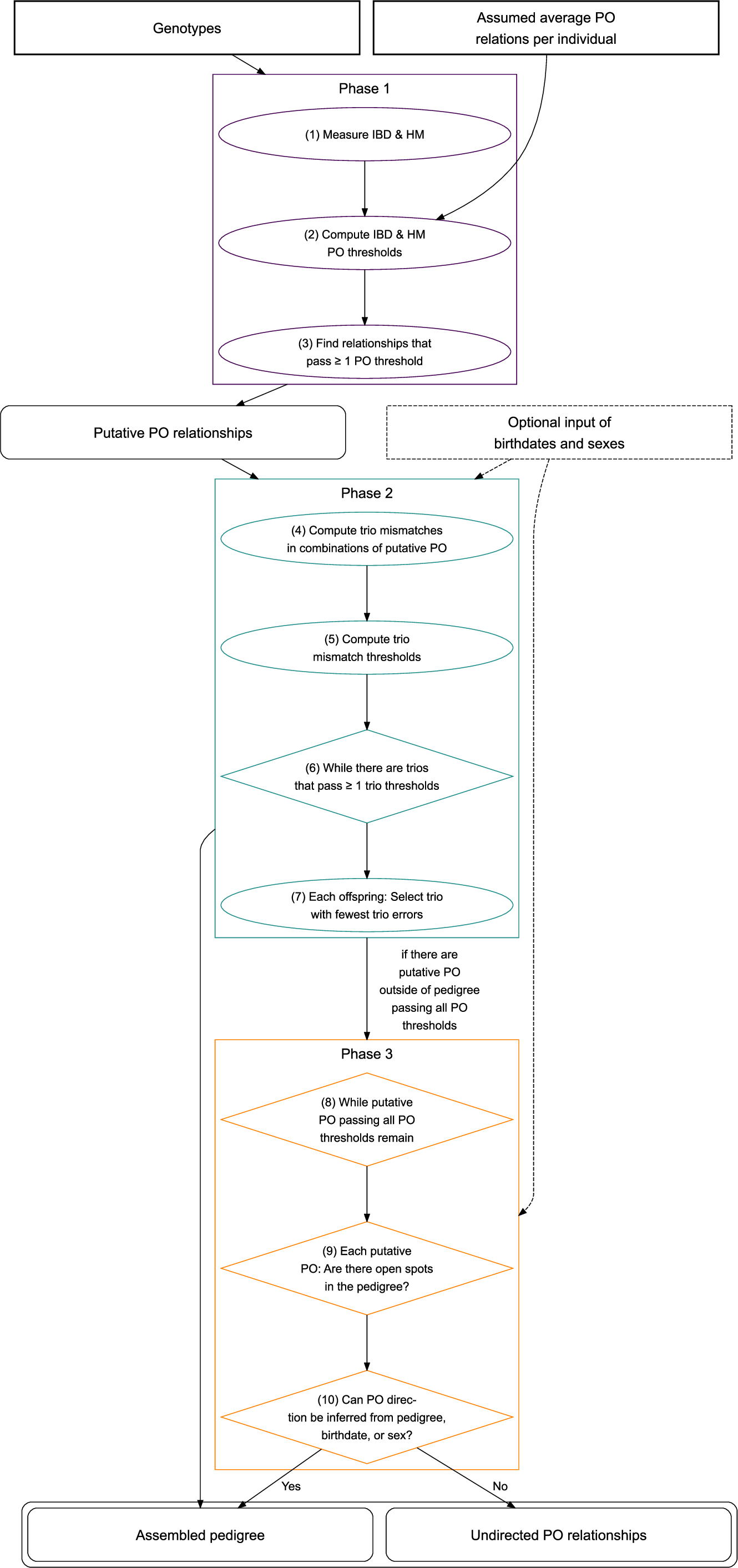
Flowchart overview of the *SPORE* algorithm. Input is shown as rectangles with square corners, while output is shown as rectangles with rounded corners. Details of each step can be found in the *Algorithm* section and in Methods.

In the first phase, *SPORE* runs *TRUFFLE* [29] to calculate IBD0 between all possible combinations of individuals. Next, *SPORE* scans a random subset of the genome to detect homozygous mismatches (opposite genotypes) between individuals, which are divided by the number of loci at which both individuals are genotyped, resulting in a relative measure of homozygous mismatches (“HM”).

Using IBD0, IBD0 IQR, and HM, *SPORE* aims to detect putative one-to-one PO relationships. To do so, *SPORE* uses the only necessary user input (apart from the genotypes themselves), the “assumed average PO relationships per individual” (APO) in the dataset (see *Performance of varying APO section* below). This input is used to automatically adjust the thresholds of IBD0, IBD0 IQR, and HM, below which a relationship is declared as putative PO.

In the second phase, *SPORE* aims to discover father-mother-offspring trios among the putative PO relationships. This phase can optionally be refined by providing birthdate and sex data. First, *SPORE* assembles putative father-mother-offspring trios out of the putative PO relationships for each focal individual that has at least two such relationships. We then quantify the number of loci that contain genotypes that could not have been transmitted from the putative parents to the putative offspring. With those Mendelian trio errors, *SPORE* once again automatically estimates the thresholds below which a PO trio is considered to be true. If multiple trios pass the threshold(s), the one with fewer errors is assigned as true.

In phase three, *SPORE* aims to infer PO relations in the remaining set of putative PO relationships that were so far not detected as a complete father-mother-offspring trio. Because these relationships can only be detected on a one-to-one rather than a more powerful trio basis, *SPORE* requires them to pass all three HM, IBD0, IBD0 IQR thresholds (see in **Figure 5**, top, how multiple thresholds decrease false positive PO detection). Furthermore, these newly inferred relationships need to fit into an empty spot of the pedigree. If provided, birthdates and sex are considered when searching for an empty spot. Once this is done for all remaining putative PO relationships, *SPORE* returns the assembled pedigree. Putative PO relationships that pass all three thresholds but could not be placed into the assembled pedigree, for example because there was no known birthdate or because both directions (parent-offspring, offspring-parent) were possible for these two individuals, are given to the user as “undirected PO relationships.”

**Figure 5:**
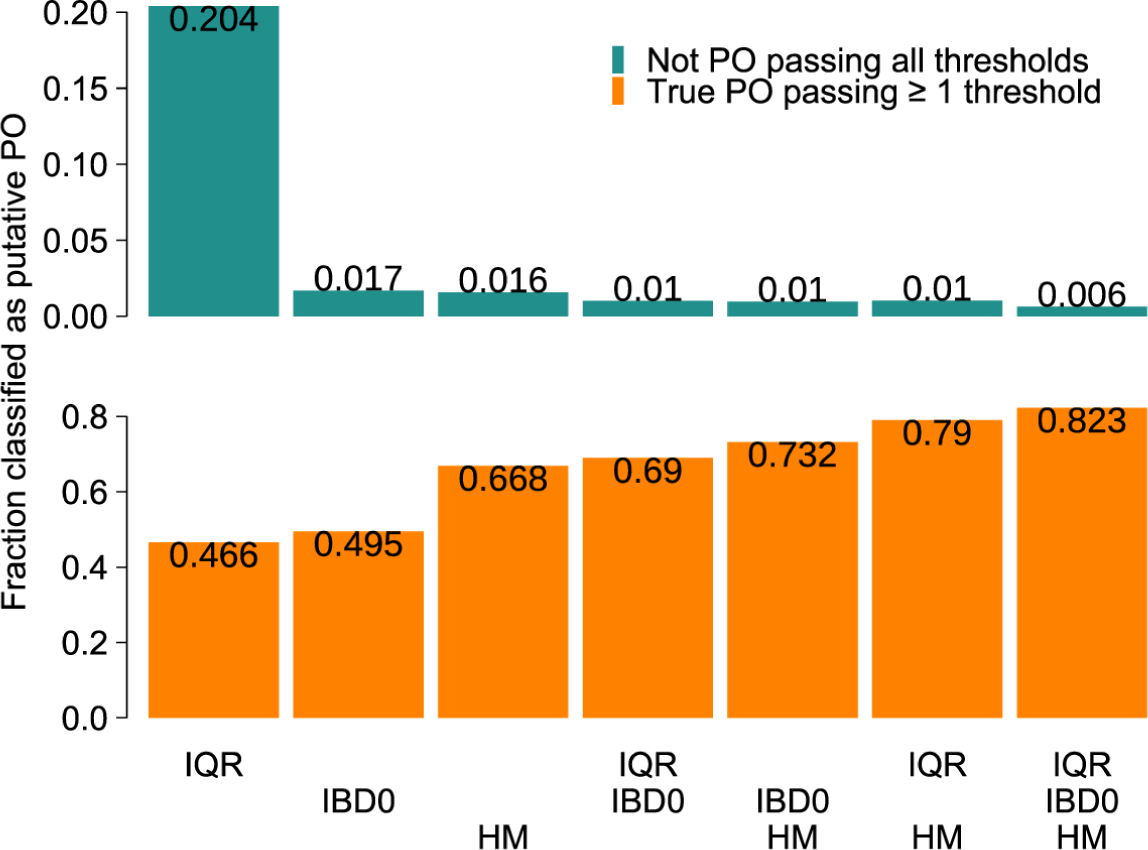
Fraction of relationships classified as putative PO in relation to which thresholds need to be passed. Top (green): fraction of relationships that are not true PO and pass all of the thresholds underneath each bar. This represents the false positives that would result from misclassification as PO in the third phase of *SPORE*. Bottom (orange): fraction of true PO relationships that pass at least one of the thresholds under each bar. This represents how well *SPORE* finds true PO in the first phase of the method. The figure is based on the simulated genotypes with APO=6.

## Comparison to other methods

We compared the performance of *SPORE* in inbred populations to *KING* [28], *CREST* [27], and *SEQUOIA* [30]. Unlike *SPORE*, these methods are not explicitly designed for inbred populations and can also infer other non-PO relationships. Like *SPORE, CREST* and *SEQUOIA* can detect the direction (i.e. who is parent, who is offspring) of relationships, while *KING* cannot. Table 1 presents an overview of these methods.

**Table 1:** Overview of the different algorithms compared with *SPORE* and some key differences between them with regards to parent-offspring inference.

| Measurement / Feature | *SPORE* | *CREST* | *KING* | *SEQUOIA* |
| --- | --- | --- | --- | --- |
| IBD | Yes, IBD0 | Yes | Yes | No |
| IBD interchrom. variation | Yes, IBD0 | No | No | No |
| Kinship | No | No | Yes | Yes |
| Homozygous mismatches | Yes | No | No | Yes |
| Mendelian trio comparisons | Yes | No | No | Yes |
| Automatic thresholds | Yes | No | No | Via likelihood |
| Infers PO direction | Yes | Yes | No | Yes |
| Birthdates input | Optional | No | No | Optional |
| Sex input | Optional | No | No | Optional |

We considered additional methods but did not analyze their performance for the following reasons. We did not further analyze *CLAPPER* [31], because it is limited to inferring relationships of individuals from five generations at a time. *PRIMUS* [19] was excluded because it did not run successfully with the IBD values in real and simulated mice analyzed here. Finally, *BONSAI* [32] was also not evaluated, because it is intended for sparse pedigrees and contains hard-coded IBD and age distributions optimized for humans.

We used three datasets to compare the methods: 1) a subset of a population of wild house mice from a long-term study, 2) five simulated datasets based on the genotypes of the founders of the long-term house mice study, and 3) a published dataset of a cattle pedigree and its genotypes [13,33].

### Zurich house mouse population

The free-living wild house mice population has been intensively monitored since 2002, when it was founded using twelve wild-caught mice from two nearby source populations [34]. The mice live in an old barn, which they can freely leave and re-enter. The barn is regularly searched for new litters and the mice are genetically sampled and genotyped at 25 microsatellite loci when they are 13 days old and again as adults (>17.5 grams of weight), and when they are found dead. The genotypes are used for identification (e.g. which pup became which adult) and pedigree construction. We analyze a random subset of 151 mice plus their parents—204 mice in total as some of the 151 mice share parents or are parents themselves—of that population, chosen at random among the individuals that we have sequenced (see *Methods*) and that have parents inferred from microsatellites. We imputed the genotypes of the mice using a custom analysis pipeline based on *AncestryHMM* [35]. A pedigree of the whole population inferred based on (incomplete) birth and death dates and (imperfect) microsatellites was only 79% complete (unpublished data), despite considerable monitoring efforts. Hence, we were motivated to explore whether genotypes derived from whole genome sequencing could also detect parent-offspring relationships and expand the current known pedigree.

### Performance

*SPORE* detected 97.7% of the true parents (*n* = 302) correctly and assigned 2 parents incorrectly when birthdates were made available and 97.7% correct with 4 wrong assignments when birthdates were not used (**Figure 6**). *CREST* did not find any parents. *KING* found 40% of parents, but also detected 92 false parents. *SEQUOIA* performed much better when it was given birthdate information. In that case, it found 32.5% of parents and only detected 3 wrong parents. Without birthdates, *SEQUOIA* found only 12.6% of parents while detecting 38 wrong parents. Hence, *SPORE* found the highest number of correct parents, while making the fewest mistakes.

**Figure 6:**
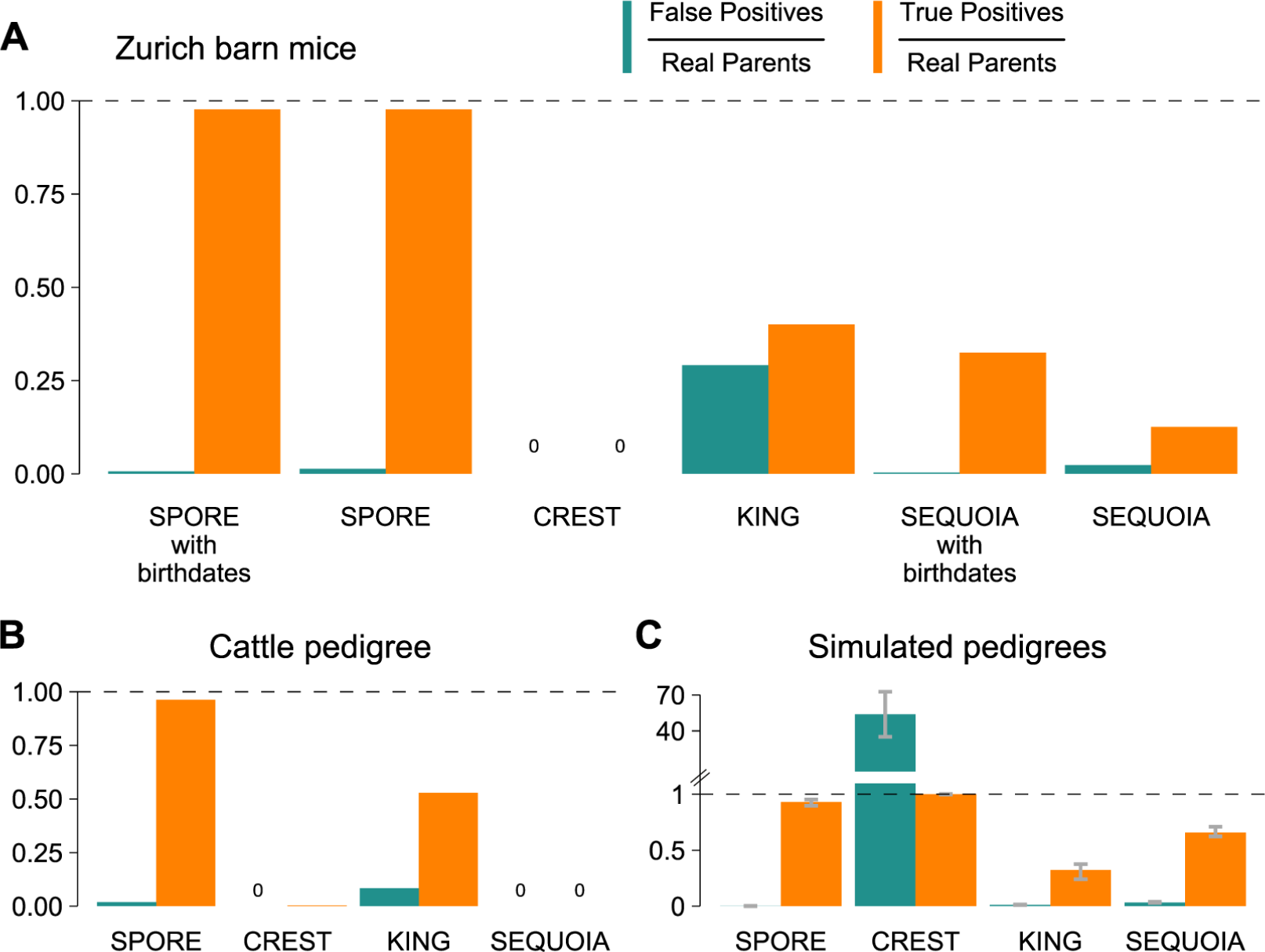
Performance of *SPORE* and three other pedigree inference methods. Two real populations are analyzed: **A)** house mice from a barn in Zurich (mean inbreeding level *F*_ROH_ = 0.3 ± 0.06) and **B)** a Holstein cattle pedigree (mean *F*_ROH_ = 0.09 ± 0.03). **C)** Performance on five simulated pedigrees (mean *F*_ROH_ = 0.37 ± 0.16). Bar heights represent the mean of each simulation and error bars denote minimum and maximum. *SPORE, CREST*, and *SEQUOIA* calls are only evaluated as true if the inferred direction of the call (who is parent, who is offspring) is correct.

### Simulated genotypes

We simulated five pedigrees (and the 19 autosomes of each individual) of randomly breeding mice of 50 overlapping generations based on the genotypes of the founders of the real house mouse population analyzed in the previous section. Due to the overlapping of generations, offspring were at most at a pedigree depth of 35 equivalents of full generations, which is the unit we refer to in the analysis of specific generations (see Methods). The pedigrees were simulated without any immigration and hence increasing levels of inbreeding.

#### Performance

*SPORE* detected 93.2% ± 2.2 (mean ± SD) of the true parents (five simulations with *n* = 4443 ± 70.5 true parents) correctly and detected on average 3.8 ± 1.8 wrong parents (**Figure 6**). *CREST* inferred all relationships to be parent-offspring, so it did find 100% of parents, but also found an average of 239, 478±67, 089 wrong ones. *KING* found only 32.3%±5.6 of parents, and also detected 52 ± 9.6 false parents. *SEQUOIA* found 65.7% ± 3.8 of parents, and 143 ± 17.3 wrong parents. Overall, *SPORE* made the fewest mistakes, while still detecting more than 90% of true parents.

We also evaluated the impact of increased inbreeding, decreased genotyping quality, and incomplete population sampling through simulated pedigrees. To test performance on inbred individuals, we analyzed pedigree generations 21-30 (**Figure 7 A**). These individuals have a considerable level of inbreeding with a mean of the genomically measured *F*_ROH_ = 0.49 ± 0.07 (see Methods). *SPORE*’s PO call accuracy in these generations was very similar to its accuracy over all generations (91% ± 3.4 vs 93.2% detected true parents with 0.08% ± 0.08 vs. 0.09% wrong parents in relation to the number of true parents). *CREST* still detected too many false parents (104, 496 ± 29, 358). *KING*’s performance was worse in these late generations than overall, detecting only 4.7% ± 4 vs 32.3% of true parents and 0.4% ± 0.4 vs 1.12% wrong parents). *SEQUOIA* was the second most reliable algorithm after *SPORE*, with 48.8% ± 8.5 parents in generations 21-30 detected vs. 65.7% parents in all generations and 5.4% ± 1 vs 3.2% wrong parents. Together, the results indicate that *SPORE* is superior at detecting PO correctly compared to other methods in controlled simulations with low and also with increasing levels of inbreeding.

**Figure 7:**
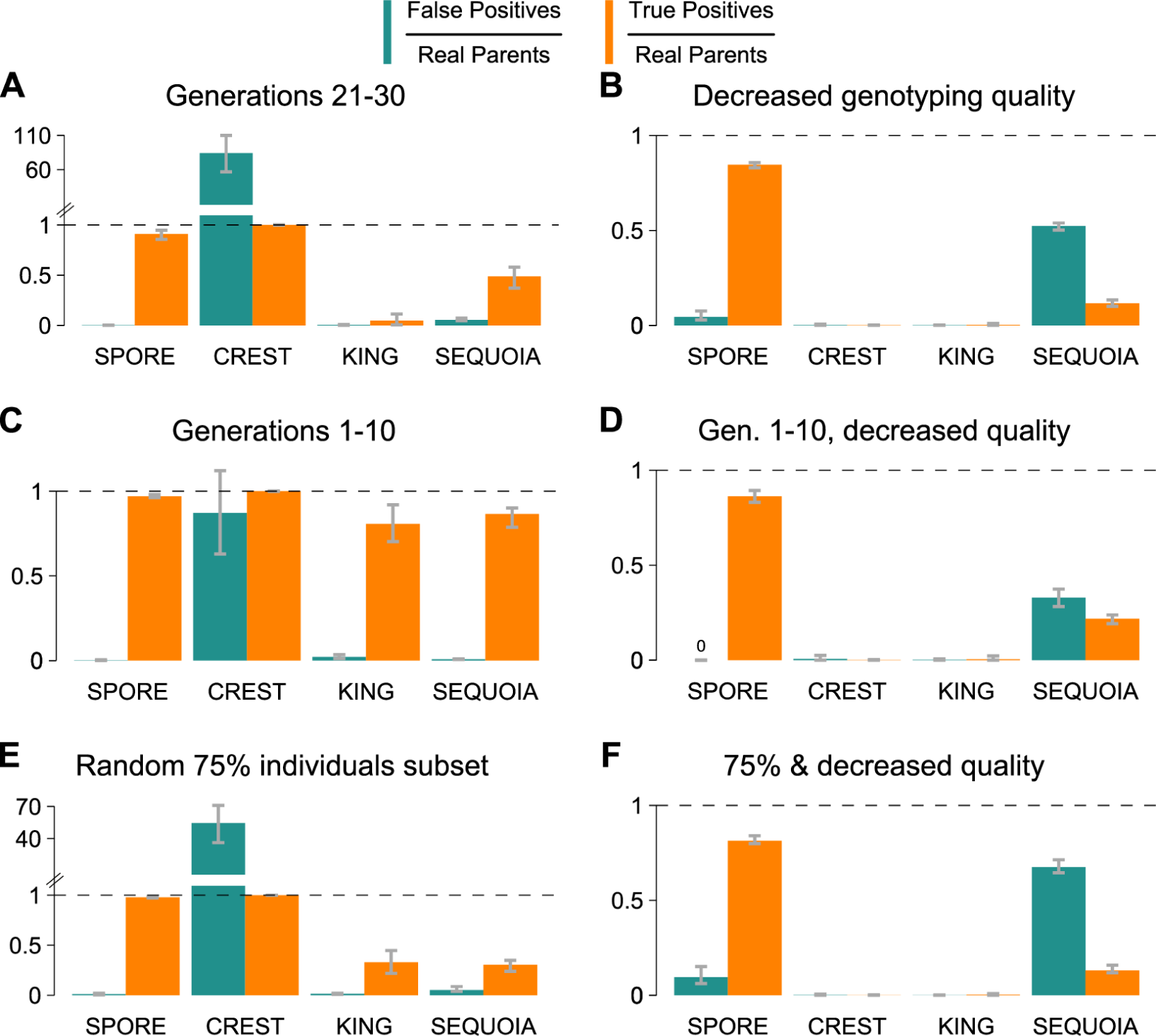
Performance of *SPORE* and three other algorithms under increased inbreeding, high genotyping errors and reduced pedigree sampling. Bar heights represent the mean of each simulation and error bars denote minimum and maximum. **A)** same simulated pedigrees as in **Figure 6 C** but only parentoffspring calls of approximately the last 10 generations are analyzed (mean *F*_ROH_ = 0.49 ± 0.07). **B)** same pedigrees as in **Figure 6 C**, but genotype imputation was performed with 10% simulated errors and 0.01x simulated coverage (rather than 2% and 0.03x). **C)** The first ten generations are analyzed (mean *F*_ROH_ = 0.17 ± 0.1). **D)** is a combination of **B** and **C**. **E)** same pedigrees and genotypes as **Figure 6 C**, but only 75% of individuals were ‘sampled’ (included in the analysis). **F)** is a combination of **B** and **E**. *SPORE, CREST*, and *SEQUOIA* calls are only evaluated as true if the inferred direction of the call (who is parent, who is offspring) is correct. *SEQUOIA* false positive calls only decrease by 5.7% (**B**) 11.4% (**D**), and 5.6% (**F**) when direction is ignored.

With decreased genotyping quality (**Figure 7 B**), but an otherwise unchanged dataset, *SPORE* still found 84.5% ± 1.2 of true parents with 4.4% ± 2 false parents. In contrast, *CREST* now only found 0.04% ± 0.07 of the true parents with 0.16% ± 0.28 false parents. Similarly, *KING* was reduced to finding 0.3% ± 0.4 of true parents and 0.05% ± 0.08 false parents. *SEQUOIA* found 11.6% ± 1.3 of true parents with 52.3% ± 1.6 false parents. Thus, *SPORE* is considerably more robust to genotyping errors than other methods.

To disentangle the effects of inbreeding and genotyping error on the inference success of the different methods, we also inferred relationships in the first ten generations (which are the least inbred at *F*_ROH_ = 0.17 ± 0.1) with and without decreased genotyping quality (**Figures 7 C,D**). *SPORE* found 97.1% ± 0.7 of true parents without decreased genotyping quality and 86.3% ± 2.4 with decreased genotyping quality, while detecting 0.1% ± 0.2 and 0% ± 0 wrong parents, respectively. In contrast, *CREST*’s results changed from detecting too many PO relationships (87.2% ± 18.2 false parents) without decreased genotyping quality to detecting almost no parents (0.06% ± 0.1 of true parents) with decreased genotyping quality. On the other hand, *KING* detected 80.7% ± 8.4 and 0.5% ± 1 of true parents with 2.1% ± 0.9 and 0.2% ± 0.3 wrong parents, respectively, showing a marked difference between genotyping qualities. Similarly, *SEQUOIA* detected 86.6% ± 4.7 of true parents with 0.8% ± 0.1 wrong parents without decreased genotyping quality, but only 21.7% ± 1.8 of true parents with 32.8% ± 3.4 false parents with decreased genotyping quality. These results are qualitatively similar to results based on individuals with *F*_ROH_ ≤ 0.05 (**SI Figure 1**). In summary, *SPORE* is more robust to genotyping errors even under minimized inbreeding.

Sampling a random 75% of individuals in the dataset (*n* = 2504 ± 83.1 true parents) and hence no longer including the entire simulated pedigree, also affected the quality of inference (**Figure 7 E**). *SPORE*, which is strongest when full trios are present in a dataset, still found 97.9% ± 0.5 of true parents, but now also 1% ± 0.74 false parents. *CREST* again detected too many false parents (136, 718 ± 36, 558). *KING* found 33% ± 9 of true parents, but also 1.5% ± 0.54 false parents. *SEQUOIA* found 30.4% ± 4 of true parents and also 5.1% ± 2 false parents. When sampling is even less complete (50%), *SPORE* still detected 93.1% ± 3.1 of PO relationships but false positives also increased to 26.9% ± 12.1 (**SI Figure 2**). *CREST* called essentially each pair as PO, while *KING* and *SEQUOIA* have a smaller increase in false positive calls than *SPORE*. Combining decreased genotyping quality with a 75% subset (**Figure 7 F**) further decreased inference quality. Nevertheless, *SPORE* still found 81.4% ± 1.8 of true parents with 9.5% ± 3.6 false parents. By comparison, *CREST* only found 0.05% ± 0.09 of true parents and 0.16% ± 0.27 false parents. *KING* detected only 0.3% ± 0.34 of true parents, and 0.06% ± 0.06 false parents. *SEQUOIA* detected 13.1 of true parents and 67.5% ± 3.3 wrong parents. Altogether, the results indicate that *SPORE* is a more accurate method when analyzing inbred, incompletely sampled pedigrees even when there are abundant genotyping errors.

##### Performance with varying APO

To evaluate the impact of the user input “assumed parent-offspring relations per individual” (APO), we set it to a range from the lowest possible (1) to quintuple (30) of the value we used otherwise (6). Overall, *SPORE* detected more true parents at higher APO, but also slightly increased the false positives (**Figure 8 A**). In the simulated pedigrees, the entire range of APO values delivered similar results, with at least 78 ± 5.5% detected true parents and at most 2.7 ± 2.2% false parents. However, APO had a bigger impact on the quality of the inference in the house mouse study population: intermediate APO values led to similar results, while extreme values led to many missed or falsely inferred parents (**Figure 8 B**). In sum, *SPORE* is not particularly sensitive to a range of APO but will have diminished accuracy at extreme values.

**Figure 8:**
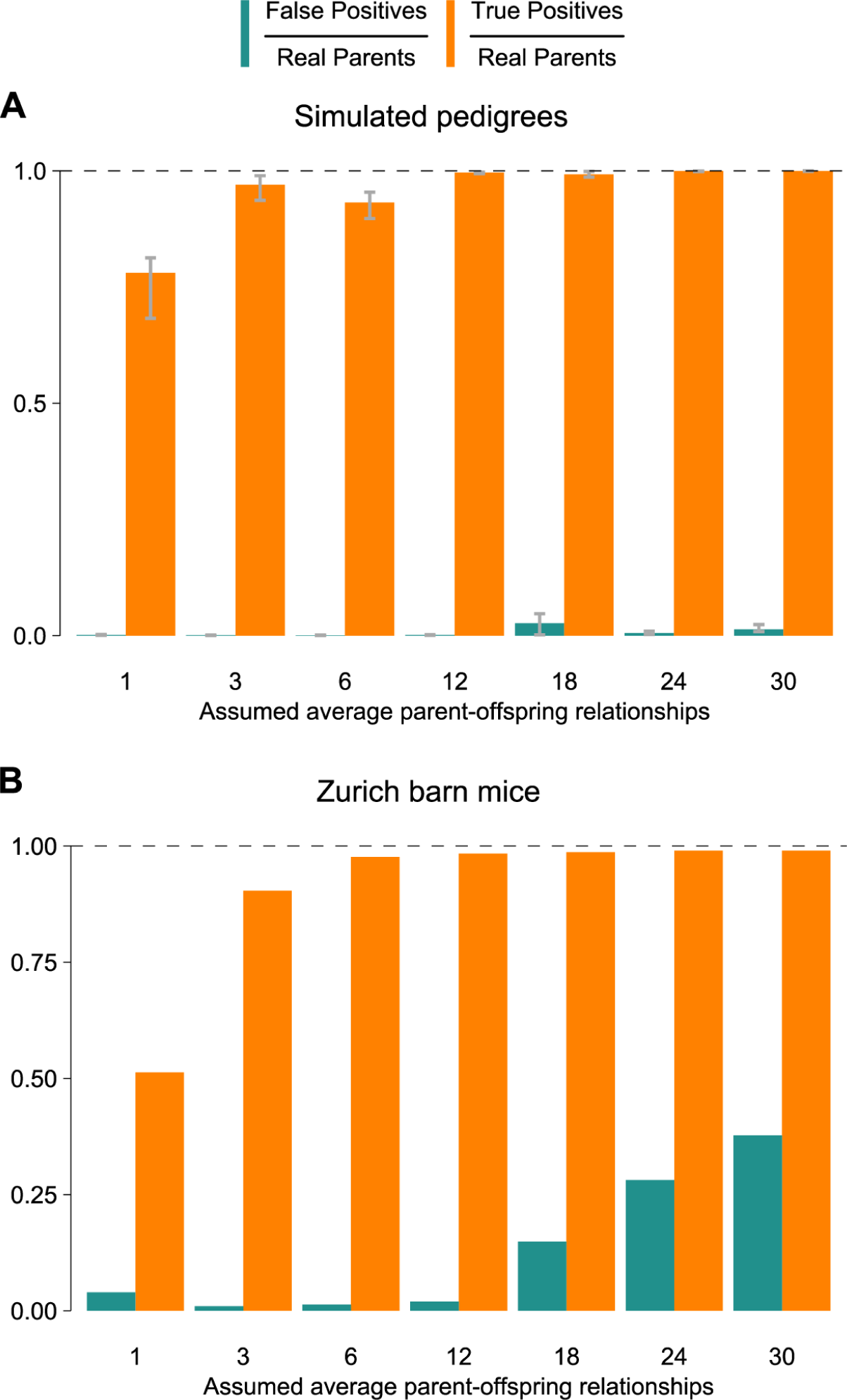
*SPORE* with different assumed average parent-offspring relationships per individual. 6 is the setting used for *SPORE* in all other plots. **A)** Bar heights represent the mean of the five simulations and error bars denote minimum and maximum. **B)** Results for the house mouse long-term study dataset.

#### Cattle population

To test *SPORE*’s performance in an independent real pedigree, we applied it to a published set of 150 cattle offspring and their 100 unique parents and some grandparents, resulting in an incomplete pedigree with 320 known PO relationships [13,33]. We found high IBD0 values between parents and offspring (mean = 0.5; SD = 0.17; expected mean and SD = 0), suggesting a high genotyping error rate that would allow us to test *SPORE* ‘s performance under these conditions.

#### Performance

*SPORE* detected 96.3% of the true parents (*n* = 320) correctly and detected 6 wrong parents (**Figure 6**). *CREST* found 0.3% of parents, with 0 false parents. *KING* found 52.8% of parents, but also detected 27 false parents. *SEQUOIA* did not find any parents. Thus, *SPORE* is superior at detecting parents in this independent real pedigree with high genotyping error rate.

## Discussion

Our results based on data from a real house mouse population, a real cattle pedigree, and simulated pedigrees, show that *SPORE* infers parent-offspring relationships in datasets with inbreeding or with high genotyping error, with considerably more accuracy than other methods.

Another advantage of *SPORE* is that it has few requirements. It does not need a genetic map and can optionally leverage sex information and birthdates. The minimal input—a VCF and a simple config file—also facilitates use by less bioinformatics-experienced investigators.

A caveat of *SPORE* is that it works on the assumption that there is at least one father-mother-offspring trio in the analyzed dataset, but this assumption is likely valid for *SPORE*’s intended uses. Furthermore, *SPORE* is designed to identify only parent-offspring relationships. However, parent-offspring inferences can then be used to identify siblings (individuals sharing parents) and to assemble multigenerational pedigrees.

We suggest that *SPORE* will be a useful tool for researchers studying experimental or wild populations with incomplete birth tracking, imperfect genotypes, and at least some autozygosity resulting from population history or recent mating between close relatives.

## Availability

*SPORE* is available at https://github.com/jnrunge/SPORE.

## Methods

Throughout different parts of the methods, we rely primarily on *R* 3.6.3 [36] with the packages *data.table* 1.13.2 [37], *dplyr* 1.0.2 [38], *ggplot2* 3.3.2 [39], *MASS* 7.3.53 [40], naturalsort 0.1.3 [41], *readr* 1.4.0 [42], *stringr* 1.4.0 [43], *sys* 3.4 [44], *tidyr* 1.1.2 [45] for *SPORE* and our analyses as well as *optiSel* 2.0.3 [46] for calculation of generations in pedigrees.

### Genomic inbreeding coefficient calculation

To calculate inbreeding coefficients in house mice, in simulated genomes, and in cattle, we used the fraction of the genomes that are part of continuous runs of homozygosity (*F*_ROH_). We used a cutoff of ≥ 1 Mb ROH [47]. Runs of homozygosity were identified using *bcftools roh* function [48], using allele frequencies of the analyzed populations. We used a fixed likelihood of 30 for unseen genotypes and a 10^-8^ M / bp genetic map, a good approximation for mammals [49]. The summed length of ROH ≥ 1 Mb on autosomes was divided by the reference length of those chromosomes, resulting in the fraction *F*_ROH_.

Values of *F*_ROH_ for other populations were extracted from the literature: Devils Hole pupfish (called using *bcftools* 1.10.2 [50], 1 Mb cutoff, extracted from plots in [16] using *WebPlotDigitizer* [51]), industry/wild chicken (called using *PLINK*, 300 kb cutoff, [14]), bottlenecked wolf (ROH identified using likelihood ratio estimation, cutoff of ≤ 10 generations recent ancestry, inferred from genetic map, [15]), Sao Paolo Quilombo (ROH detected with *PLINK*, 500 kb cutoff, [25]), and UK Biobank (called with *PLINK*, no cutoff, [26]).

### *SPORE* (Specific Parent-Offspring Relationship Estimation)

#### Speed

*SPORE* runs for 8 hours on one core for a pedigree of 2500 individuals. However, *SPORE* can take advantage of multiple cores in some key parts of its algorithm, approximately halving its run time on four cores compared to one and *SPORE* can also optionally skip steps when rerunning it on the same data. Details on the speed of *SPORE* can be found in **SI Figures 3 & 4**.

#### IBD calculation

We used *TRUFFLE* 1.38 [29] to calculate IBD0 values for each relationship. We excluded variants with a minor allele frequency `-maf` below 10^-4^ and/or >95% missing genotypes at that locus. Both settings can be customized in the *SPORE* options file. To extract IBD0 values for all chromosomes separately, we also ran *TRUFFLE* on VCF files containing only one chromosome at a time. This is automatically done within *SPORE* for all chromosomes found in the input VCF file. IBD0 IQR was calculated as the interquartile range—which is less sensitive to outliers than the variance—of all IBD0 values of each of the autosomes.

#### Homozygous mismatches

We calculate homozygous mismatches on a user-specified random fraction of the genome to increase speed. The *SPORE* default and value used in this manuscript is 0.01. Impacts of other values on speed and results are shown in **SI Figures 4-6**. We then run a custom *R* script that counts the loci with homozygous mismatches for every pair of individuals and the loci where both individuals compared are genotyped. We then calculate the fraction of loci genotyped in both individuals that have homozygous mismatches for each pair.

#### Mendelian trio mismatches

We calculate genotype mismatches in possible father-mother-offspring trios using the *mendelian* plugin of *bcftools*. Trios are deemed possible and thus worthy of this time-intensive computation in all cases where a given individual (the putative offspring) 1) has a putative PO relationship to both putative parents, 2) optionally available birthdates are not older for the offspring than for the parent, and 3) optionally available sexes differ between the two parents. All trios fulfilling these criteria are then passed on to *bcftools* for the computation of the mismatches. We extract a relative measure of these mismatches by dividing the number of loci with mismatches by the number of loci that were genotyped in all three individuals in a putative trio. A mismatch can be a heterozygous offspring genotype when both parents are homozygous for the same allele or a homozygous offspring genotype when each parent is homozygous for opposite alleles.

#### Thresholding

##### Putative PO relationships

To calculate thresholds below which relationships are considered as putative PO, *SPORE* calculates the number of individuals that have ≥ 1 PO relationship and > APO PO relationships at levels of IBD0, IBD0 IQR, and HM, separately. This is tested in up to 10,000 increments from the lowest value to the highest, with the first 9,000 increments being exactly the 9,000 lowest values, and the last 1,000 increments being equally sized from the 9,000th lowest value to the highest value (**SI Figure 7 A**). Fewer than 10,000 increments are used if fewer than 10,000 unique values exist of the variable that is looked at. The greatest distance between the fraction of individuals having at least one PO relationship and those that have more than the APO number of PO relationships is then set as the threshold of that variable below which relationships are classified as putative PO. At this stage, passing one threshold is enough to be classified as a putative PO relationship (see **Figure 5**, orange, to see how multiple thresholds improve PO detection).

##### Putative father-mother-offspring trios

To determine the threshold below which Mendelian trio errors are low enough to consider the father-mother-offspring trio as true, Mendelian errors are used in two ways. In the first case, we aim to quantify how a trio’s Mendelian errors compare to other trios that were evaluated for the same offspring. To that end, a trio’s Mendelian errors are divided by the mean Mendelian trio errors of the other evaluated trios with the same putative offspring (“offspring-relative trio errors”). This could be useful if genotype quality varies between individuals, which could increase the errors in both the true and false trios. In the second case, we seek to analyze how low a trio’s Mendelian errors are compared to all evaluated trios (of all putative offspring) across the dataset. The resulting variable is simply the percentile that each relative error count falls in compared to *all* relative error counts (“dataset-relative trio errors”). We calculate this using the `ecdf` function of *R*.

For both offspring- and dataset-relative trio errors, *SPORE* then estimates the thresholds below which a trio is considered as true. Since only one set of parents can be true per offspring and only trios made up of putative PO relationships are tested, *SPORE* makes use of two values to estimate the thresholds, 1) the fraction of putative offspring that would have exactly one trio, and 2) the fraction of putative offspring that would have more than one trio at each threshold (**SI Figure 7 B**). This is tested in up to 10,000 increments from the lowest value to the highest, with the first 9,000 increments being exactly the 9,000 lowest values, and the last 1,000 increments being equally sized from the 9,000th lowest value to the highest value. Fewer than 10,000 increments are used if fewer than 10,000 unique values exist of the variable being analyzed. The threshold is then set at the trio errors value at which the highest fraction of putative offspring have exactly one trio that falls below the threshold, but this fraction cannot be higher than the fraction of putative offspring that have more than one trio that passes the threshold. If only one trio was evaluated for a given putative offspring, only the dataset-relative trio error threshold is used. In case more than one trio passes a threshold for a putative offspring, i.e. two sets of parents seem true, *SPORE* chooses the trio that has fewer errors. These cases are also output separately for the user to have the opportunity to examine more closely.

### Execution of other algorithms

*CREST* 1.0.0 was run on IBD data inferred by *IBIS* 1.20.9 [52], as recommended by *CREST* developers [27]. *IBIS* was run using the genetic map of the Collaborative Cross from [53] for *Mus musculus* and simulations and the genetic map from [54] for cattle.

*KING* 2.2.5 had to be used with pruned genotypes (using *PLINK*) as it would otherwise not complete. Parameters were chosen to prune as little as necessary for execution to succeed. In the end, we chose 200 kb windows that have a correlation of *R*^2^ < 0.3 for the long-term study, *R*^2^ < 0.5 in 250 kb windows for the simulations, and *R*^2^ < 0.75 in 10 kb windows for cattle. There was one exception in the simulations that required pruning in 300 kb windows in the “decreased genotyping quality” condition for *KING* to successfully complete analysis.

*SEQUOIA* 1.3.3 was run without sibling inference and both pedigree parent-offspring relationships and potential parent-offspring relationships (`GetMaybeRel` function) were considered, though the latter gave only few additional results. We pruned the genotype data because *SEQUOIA* is designed to run on genotypes in the hundreds. We used *R*^2^ < 0.1 in 1 Mb windows for the long-term study population of house mice and simulated genotypes, with the exception of “decreased genotyping quality” conditions, which required pruning to *R*^2^ < 0.01 in 5 Mb windows to allow *SEQUOIA* to finish running (within 5 days). We used *R*^2^ < 0.01 in 10 Mb windows for the cattle due to the larger number of loci in the unpruned dataset (ten times more than in the house mouse datasets).

Sex information was provided to *SPORE* and *SEQUOIA* in all runs, while birthdates were only made available to both in the long-term mouse data.

### Datasets

#### Zurich house mouse population

##### Sequencing

We sequenced the twelve founders (*F*_0_) of the population and eight of their *F*_1_ offspring (at least one for each founder) to an average of 9.2x (SD=1.16) coverage after removing duplicates (reads available in SRA under BioProject PRJNA782421). We used the Illumina HiSeq X platform with paired-end reads of 150 base pairs. We aligned all reads to the reference *Mus musculus* genome (GRCm38.p6) using the *mem* algorithm of bwa 0.7.17-r1188 [55], sorted the resulting bam files and marked duplicates using Picard toolkit 2.18.26 [56]. We called genotypes using *bcftools* 1.10.2 [50] and *Strelka* 2.9.10-0 [57], both filtered independently to only include autosomal bi-allelic loci with a minimum variant quality (‘QUAL’) of 100, no missing genotypes, ≥ 3 reads per genotype, but otherwise no genotype quality filtering at this stage. We then used the *isec* function of *bcftools* to extract those genotypes that were supported by both genotype callers. Next, we removed loci with genotypes that were found to disagree within the *F*_0_-*F*_0_-*F*_1_ trios (Mendelian errors) using *vcftools* 0.1.16 [58]. Finally, we removed loci that were found to be in regions of particularly high (> 25x) or low (< 5x) average coverage across the *F*_0_*F*_1_. We phased those genotypes using the known pedigree with *WhatsHap* 0.18 [59] based on the sex-averaged genetic map published by [53]. Afterwards, we only kept loci in which all genotypes had at least a genotype quality score of 20. In the end, we detected 2,161,810 loci with variation among the founders.

We then sequenced 204 individuals (making up the 302 unique parent calls) from later generations, *F_x_*, on the Illumina NextSeq 500 platform with single-end 75 base-pair reads and alternatively on the Illumina NovaSeq platform with paired-end 150 base pair reads to an average 0.03x (SD=0.01) effective coverage (the fraction of loci of interest with at least one read, see [60]). We aligned all reads to GRCm38.p6 using the *mem* algorithm of bwa and marked duplicates using Picard. From these aligned reads, we extracted positions at the 2,161,810 loci with variation in the founders with a base quality ≥ 30 using the *mpileup* function of *samtools* 1.10 [61]. For each individual, we excluded variants within 300 bases of another variant to reduce bias towards one strand due to several bases being in one read pair. These bases were then used as input for genotype imputation.

##### Genotyping via imputation

To genotype the house mice in the long-term study, we designed an imputation pipeline that converts the output of the ancestry inference software *AncestryHMM* 0.94 [35] into genotypes. In short, the pipeline takes phased genotypes of the twelve founding mice of the population and uses these as 24 ancestral genomes. From these ancestral genomes and the low-coverage data of each of the 204 individuals that we analyzed, *AncestryHMM* infers probabilities for each ancestry combination at each locus. Each of the 24 ancestral genomes was assumed as having equal representation in the overall population, and for each individual the foundation of the population was assumed to be 10 generations in the past, but this was not fixed, i.e. *AncestryHMM* could deviate from it. We assumed a genotype error rate of 0.1% and used a genetic map with 10^-8^ M / bp.

*AncestryHMM* outputs the probability of each ancestry combination (e.g. ancestry 1 and 3) at each locus. To convert this into genotypes, we first interpolated the probabilities at loci in between the ones where bases were available. Next, we summarized the probabilities for each ancestry by taking the probability for this locus to be homozygous for ancestry *y* and adding half of the sum of all probabilities for this locus to be heterozygous for ancestry *y*, because the probability to be heterozygous represents the probability for only one chromosome (per locus) to descend from this ancestry.

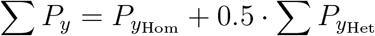

We multiplied this total probability *P_y_* for ancestry *y* at each locus with the allele of this ancestry (reference allele = 0.0, unphased = 0.5, and alternative allele = 1.0). We did this for all 24 ancestries and then summed these allele values up at each locus to receive an imputed proportion of the alternative allele for each locus of each offspring (between 0 and 1). Imputed proportions between 0.0 and 0.01 were inferred as a homozygous reference allele genotype, between 0.495 and 0.505 as heterozygous, and between 0.99 and 1.0 as homozygous alternative allele. This means that 99% of the *AncestryHMM* inferred ancestries agreed on the genotype.

For the long-term study population of house mice, we removed genotypes in the first and last 10 Mb of each chromosome for the analyses here, because we detected a slightly increased genotype error rate in those regions.

### Simulated pedigrees

We simulated genotypes of mice for 50 time intervals (~ generations, which can overlap) using *SimuPOP* [62] based on the genotypes of the 12 mice that founded the long-term study population, the same that were used as ancestral alleles for imputation. Genotypes were simulated at their respective loci, i.e. loci kept the same position as implied by the reference genome (GRCm38.p6). The genetic distance in Morgans between loci, and thus the probability of recombination, was based on the sex-averaged map published by [53].

We simulated overlapping generations in which each individual lives for two time intervals after their birth and could mate in both of those. Individuals were simulated to mate randomly, but could only produce Pois_50_ offspring each generation, and each randomly mated pair would produce Pois_6_ offspring, with Pois_λ_ being a Poisson distribution. Hence, by chance, some individuals could have multiple litters with Pois_6_ offspring, but many would also have no offspring in a generation. Litter size and offspring per time interval are within the typical ranges observed also in the long-term study [63,64]. We simulated five unique pedigrees with these settings.

To simulate erroneous and low-coverage sequencing, we extracted bases from all the simulated genotypes of each simulated individual with a probability of Pois_cov_, with “cov” being the coverage (0.03, with the exception of 0.01 in “decreased genotyping quality” analyses). Furthermore, each extracted base was simulated to be ‘misread’ as the other possible base from its bi-allelic locus with a probability of Pois_err_, with “err” being 0.02 or 0.1, with 0.02 being used in the main analyses and 0.1 in the “decreased genotyping quality” analyses.

The simulated reads were then used as the basis for the imputation pipeline described above. Hence, all analyses of simulated mouse genotypes presented here are based on imputed genotypes, which contain some errors due to either limits of the imputation in general or the base errors that we deliberately introduced. This is done to create a dataset that is comparable to the genotypes of the real house mice we also analyzed.

In the end, the simulated pedigrees cover a wide range of inbreeding levels, measured through *F*_ROH_ (**SI Figure 8**). Any mention of the generation of an individual is based on the ‘true’ pedigrees of the simulations and the summary statistic “equiGen” of the *R* package *optiSel*, which represents the “number of equivalent complete generations,” i.e. is adjusted for matings within generations. When only specific generations were analyzed, the output files of the methods and the true pedigree were subset to only include relationships and trios within individuals from the targeted generations, e.g. if one parent was not of the targeted generations, none of their children would be included in the analyses.

### Cattle

We converted the published set of 13,037,955 loci (derived from whole-genome short-read Illumina sequencing using Illumina HiSeq 2000 by [33]) into VCF format. We filtered out loci with more than 10% missing genotypes and loci without allelic variation, and only kept the autosomes for the analyses, yielding a final set of genotypes at 12,653,341 loci.

## Ethics

The data were collected under permits 26/2002, 210/2003, 215/2006, 51/2010 from the Swiss Animal Experimentation Commission.

## Author contributions

**Conceptualization:** JNR, AL, BK, AB. **Investigation:** JNR, AL. **Formal analysis:** JNR. **Methodology:** JNR, AB. **Resources:** AL, BK. **Software:** JNR. **Visualization:** JNR. **Supervision:** AB. **Funding Acquisition:** JNR, AL, BK, AB. **Writing:** JNR, AL, BK, AB.

## Acknowledgements

Jari Garbely extracted DNA of house mouse samples and Kerel Francis prepared sequencing libraries. Sally Steinert, Bruce Boatman, and many others helped monitor and sample the mice in the barn of the long-term study. Itsik Pe’er provided comments on the manuscript.

## Data availability

Sequence reads of the house mice from the long-term study available in NCBI Sequence Read Archive under BioProject PRJNA782421 at acceptance.

