## Supplemental Figures for "Parent-offspring inference in inbred populations"

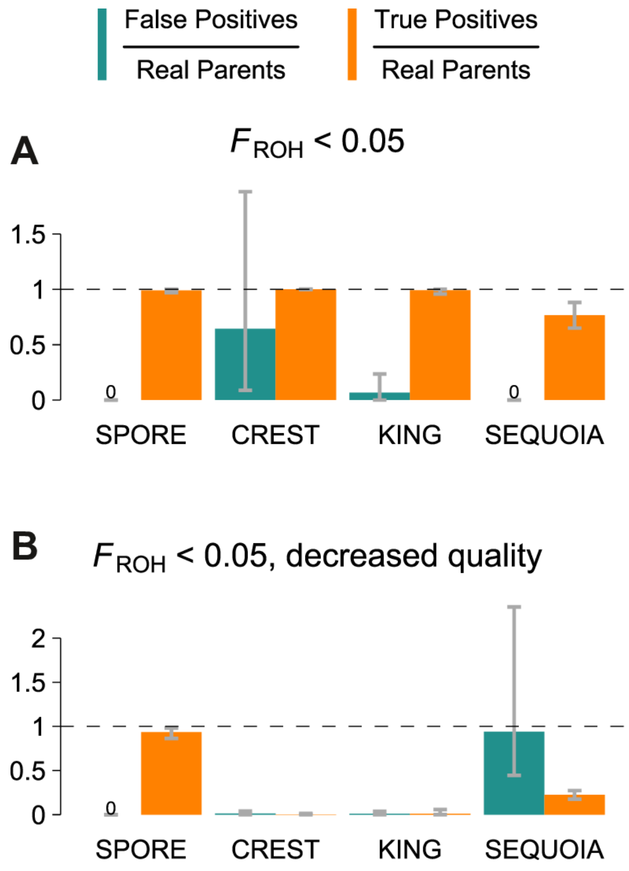


**Suppl. Fig. 1: Parent-offspring inference performance with minimal inbreeding.** Bars indicate the mean of five simulated pedigrees and the error bars show minimum and maximum values. These plots only include calls on individuals with $F_{\text{ROH}} \leq 0.05$, which is only the case for $51 \pm 25$ parent-offspring relationships. **A)** Simulations with standard amount of genotyping errors. **B)** Simulations with increased genotyping errors. *SPORE*, *CREST*, and *SEQUOIA* calls are only evaluated as true if the inferred direction of the call (who is parent, who is offspring) is correct. *SEQUOIA* false positive calls only decrease by 3.4% (**B**) when direction is ignored.

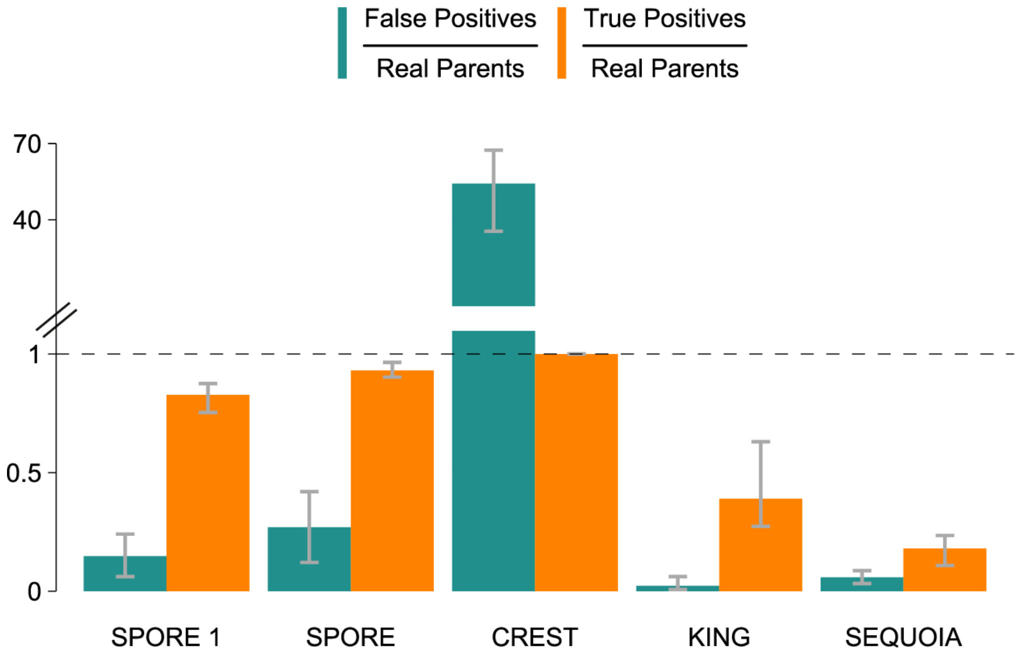


**Suppl. Fig. 2: Parent-offspring inference performance in pedigrees were only 50% of the population was sampled (at random).** Bars indicate the mean of five simulated pedigrees and the error bars show minimum and maximum values. "*SPORE 1*" indicates results based on APO=1, while "*SPORE*" uses APO=6 as in all other plots. *SPORE*, *CREST*, and *SEQUOIA* calls are only evaluated as true if the inferred direction of the call (who is parent, who is offspring) is correct.

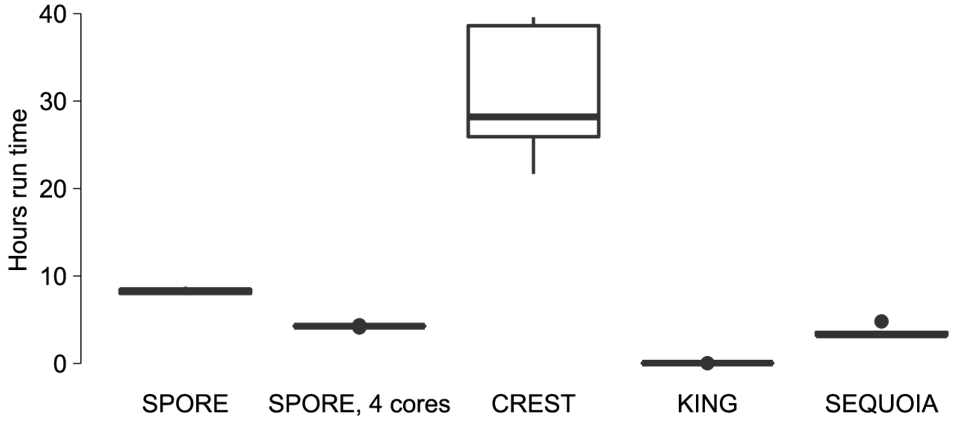


**Suppl. Fig. 3: Runtime length in the simulated data sets for the four algorithms.** For a practical comparison, these times include IBD detection (*TRUFFLE* for *SPORE* and *IBIS*, as recommended, for *CREST* ) and pruning (*KING*, *SEQUOIA*).

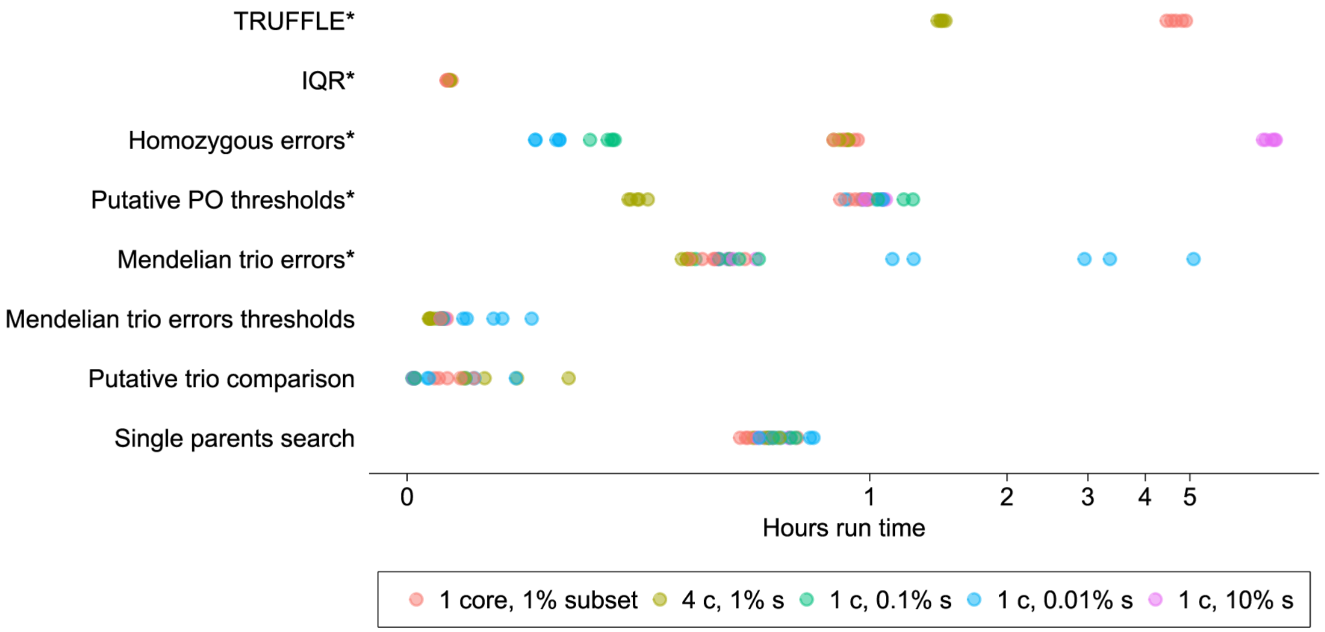


**Suppl. Fig. 4: Runtime length in the simulated data sets for different steps of *SPORE*.** Colors indicate differently sized random subsets of the input variants to calculate homozygous mismatches and number of cores used (purple). Steps marked with an asterisk can optionally be skipped by providing/keeping the output from a previous *SPORE* run, for example if one only wants to change the APO value.

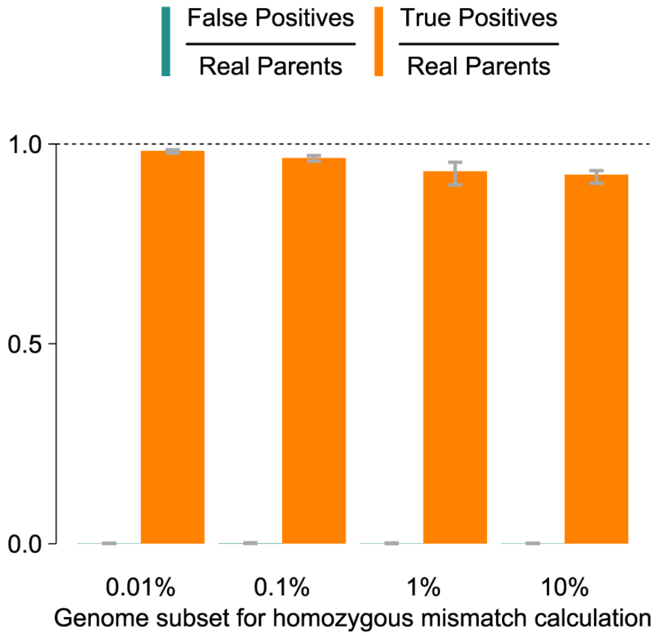


**Suppl. Fig. 5: Varying genome subsets as a basis for calculation of homozygous mismatches in simulated pedigrees.** Bars indicate the mean of five simulated pedigrees and the error bars show minimum and maximum values. Smaller values appear to improve the results, likely due to more individuals being included in putative trio comparisons. This is helpful when datasets are complete and the correct parents can be found.

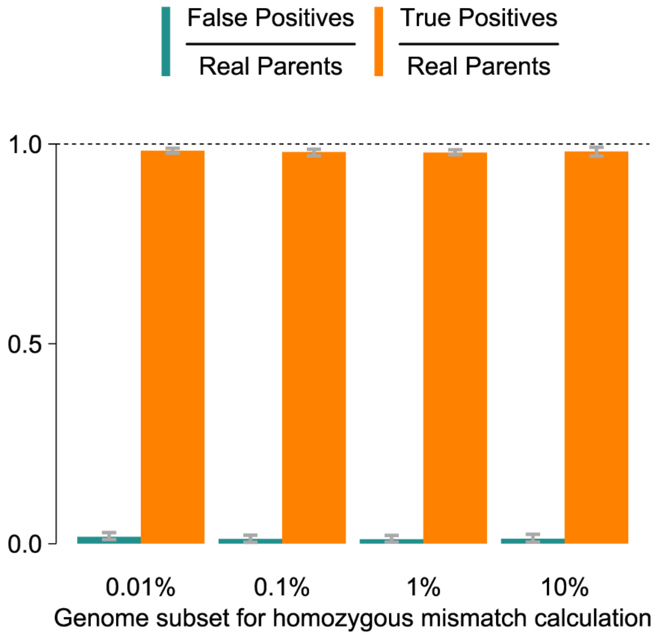


**Suppl. Fig. 6: Varying genome subsets as a basis for calculation of homozygous mismatches in simulated pedigrees, randomly subset to 75% of individuals.** Bars indicate the mean of five simulated pedigrees and the error bars show minimum and maximum values. In contrast to **SI Figure 3** above, smaller values no longer improve results, because suboptimal pre-selection of putative PO will now lead to the inclusion of more wrong relationships, which impacts the quality of the chosen trios.

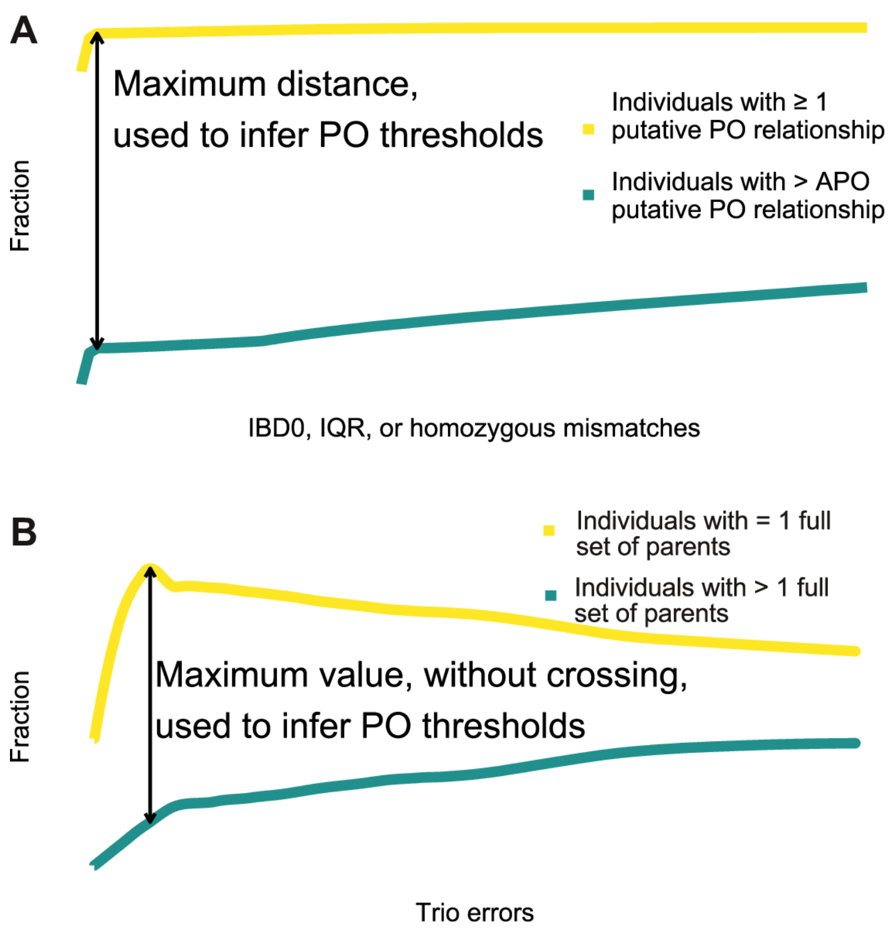


**Suppl. Fig. 7: Example plots showing how automatic threshold calculation in *SPORE* works.** The illustrations are based on averaged simulation data. **A)** Automatic thresholding for putative PO. In yellow, the fraction of individuals with $\geq 1$ putative PO relationship if the x axis value was the threshold below which relationships are classified as PO. In green, the fraction of individuals with more PO relationships than the "assumed average PO relationships per individual," APO. A distance-drawing line highlights what *SPORE* selects as the threshold, the maximum distance between the two fractions. **B)** Automatic thresholding for trios. In yellow, the fraction of individuals that has exactly one full trio at the x-axis threshold. In green, the fraction of individuals that has more than one full trio. The threshold is then set to be the highest point on yellow as long as green has not crossed yellow.

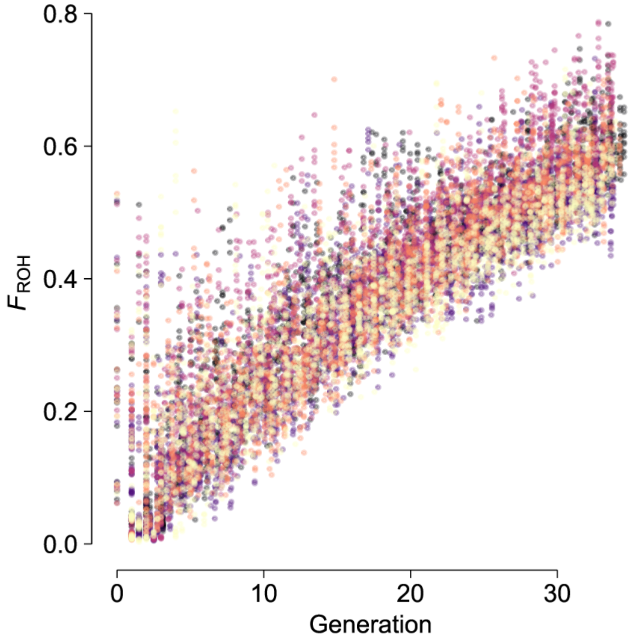


**Suppl. Fig. 8: Individual inbreeding $F_{\mathbf{ROH}}$ values for all simulated individuals.** The x axis represents the generation based on the true pedigree of the simulations. Colors denote the five unique simulated pedigrees.
